# Two simple criteria to estimate an objective’s performance when imaging in non design tissue clearing solutions

**DOI:** 10.1101/799460

**Authors:** Sabrina Asteriti, Valeria Ricci, Lorenzo Cangiano

**Affiliations:** Dept. of Translational Research, University of Pisa, Pisa, Italy; Dept. of Neurosciences, Biomedicine and Movement Sciences, University of Verona, Verona, Italy; Dept. of Mathematics and Computer Science, University of Palermo, Palermo, Italy

**Keywords:** fluorescence, microscopy, tissue clearing, spherical aberration, serial optical sectioning, deconvolution

## Abstract

Tissue clearing techniques are undergoing a renaissance motivated by the need to image fluorescence deep in biological samples without physical sectioning. Optical transparency is achieved by equilibrating tissues with high refractive index (RI) solutions, which require expensive optimized objectives to avoid aberrations. One may thus need to assess whether an available objective is suitable for a specific clearing solution, or the impact on imaging of small mismatches between cleared sample and objective design RIs. We derived closed form approximations for image quality degradation versus RI mismatch and other parameters available to the microscopist. We validated them with computed (and experimentally confirmed) aberrated point spread functions, and by imaging fluorescent neurons in high RI solutions. Crucially, we propose two simple numerical criteria to establish: (i) the degradation in image quality (brightness and resolution) from optimal conditions of any clearing solution/objective combination; (ii) which objective, among several, achieves the highest resolution in a given immersion medium. These criteria apply directly to the widefield fluorescent microscope but are also closely relevant to more advanced microscopes.

## 1 INTRODUCTION

The traditional approach to the imaging and 3D reconstruction of biological tissue involves serial or blockface physical sectioning to obtain a closely-spaced sequence of 2D images, followed by processing to merge these planar datasets into a volumetric representation. However, the development of transgenic animals expressing fluorescent proteins linked to specific promoters (e.g. [1]) has led to a shift toward non-destructive imaging of intact and cleared samples. In this approach tissues, including entire organs, are made optically transparent by reducing their refractive index (RI) inhomogeneities [2]. In practice, interstitial and intracellular water is replaced with a high RI solution, optionally combined with the chemical removal of lipid scatterers. Thus, any location within the volume of a thick sample can be viewed simply by adjusting the microscope’s *object plane* by focusing, and the entire sample can be imaged through a process of serial optical sectioning. While optical sectioning is more time efficient than physical sectioning, it has several limitations.

First, light emitted by sources located above and below the object plane enters the objective and reaches the detector, which reduces the signal-to-noise ratio. Laser-scanning confocal and light sheet microscopes limit this phenomenon to some degree compared to the common widefield fluorescent microscope. In all cases, a further improvement can be obtained *a posteriori* by the computational operation of deconvolution [3]. A luminous point source is viewed at the camera as a complex 2D pattern called the *point spread function* (PSF), determined by diffraction by the objective’s aperture (and other parameters in more advanced microscopes). In deconvolution, knowledge of the system’s set of 2D PSFs as a function of focus position (the so-called ‘3D PSF’), enables to reassign out of focus light to its location of origin in the sample.

A second limitation is linked to the fact that different clearing protocols equilibrate tissue with solutions having different RIs. Ideally, the objective used for imaging would be designed for the exact RI of the chosen clearing solution. In reality, mismatches between the objective design and tissue clearing solutions introduce an additional perturbation in the form of aberration, leading to a more extended PSF [4–8]. The consequences, often dramatic, are an increase in out-of-focus light and a decrease in spatial resolution. Deconvolution with an appropriately expanded PSF can restore image quality only up to a point, due to the irreversible loss of spatial frequency information (the so-called ‘missing cone’ problem; [9, 10]).

Clearly, the selection of an appropriate combination of clearing medium and objective is critical to avoid severe degradation of image quality. Many papers rigorously characterized the aberrations introduced by RI mismatches, but few easy-to-apply guidelines are available to the microscopist [11]. Here we propose two numerical criteria that are simple to calculate and only use standard parameters. They model an objective immersed in its design solution and separated from the cleared sample by a coverslip, or the specific case of the objective immersed in a mismatched clearing solution.

## 2 MATERIALS AND METHODS

### 2.1 Optics

The imaging system consisted of a DM LFSA upright microscope (Leica Microsystems, Wetzlar, Germany) equipped with a 49020 narrow band EGFP filter set (Chroma, Bellows Falls VT, USA) and a DFC 350FX cooled monochrome 12 bits CCD camera (Leica) coupled with a 0.63x tube. Image stacks were acquired with µManager software [12]. The objectives used for imaging or modeling were water immersion for electrophysiology (i.e. long working distance) from Leica Microsystems: 0.30 NA (#15506142, 10x, WD_*d*_=3.60 mm), 0.50 NA (#15506147, 20x, WD_*d*_=3.50), 0.80 NA (#15506155, 40x, WD_*d*_=3.30).

### 2.2 Image processing environment

Numerical integration of the modified Gibson & Lanni (G&L) model and all image analyses were performed with Fiji/ImageJ [13, 14] using public domain plugins, open source and custom scripts.

### 2.3 Measurement of the experimental PSF

Green PS-Speck fluorescent microspheres (P7220; Thermofisher Scientific) with diameter 175 nm (SD 5 nm) were diluted 1:1000 with 1% w/v low gelling temperature agarose (A9414; Sigma-Aldrich/Merck) solution at 37 ℃ and vortexed. The suspension was polymerized in a 2–3 mm thick convex meniscus on the bottom of a Petri dish covered with black filter paper (AABP02500; Sigma-Aldrich/Merck). The top of the gel meniscus was removed with a vibratome (VT 1200S; Leica Biosystems) to obtain an optically flat surface, covered with 20% FRUIT clearing solution [15] and agitated continuously. The solution was replaced regularly until the RI of the gel matched that of 20% FRUIT. RI was measured with a calibrated refractometer (ORA 4RR; Kern-Sohn). All procedures were performed in far red light or darkness. Before acquisition the Hg lamp was allowed to stabilize for >30 min.

A volume was acquired with the 0.5 NA objective as a stack of 201 slices (1000 ms exposure/slice) at sampling intervals of 325×325×500 nm/voxel, somewhat below (in x-y) and well beyond (in z) the Nyquist sampling interval in diffraction-limited conditions. The stack was converted to 32 bits and all well-isolated diffraction patterns extracted, upsampled, aligned and averaged together with PSF Creator and PSF Combiner in the GDSC-SMLM suite by Alex Herbert (University of Sussex). The output, a single stack containing the 3D measured PSF, was then converted to a 2D axial section (Figure 4), as follows: (i) the axis of symmetry of the PSF was determined precisely; (ii) for each slice at axial coordinate *Z*,all pixels overlapping a circle of radius *R* centered on the optical axis were averaged together.

### 2.4 Simulation of the model PSF with sampling by the CCD

A 3D PSF was generated with the modified G&L model for the 0.5 NA objective at a 16 fold x-y sampling rate relative to that of the CCD (20×20×500 nm/voxel). Multiple copies of this PSF, each shifted in x and/or y by 0, 4, 8, 12 pixels, were assembled in a combined stack and downsampled by 16 fold in x-y by binning. The resulting stack was further processed in the same way as the experimental one containing fluorescent microspheres (section 2.3).

### 2.5 Imaging of fluorescent neurons in design and mismatched solutions

The mouse line used and the spinal cord dissection procedures were described previously [16]. Briefly, an early postnatal (P4) Galanin-eGFP^+/+^ [17] male mouse was sacrificed with approval by the ethical committee of the University of Pisa (n. 10/2018 as per D.lgs.vo 26/2014) and in accordance with EU Directive 2010/63/EU. The spinal cord was dissected and fixed with 4% paraformaldehyde in phosphate buffered saline (PBS), laid on black filter paper and covered with 1.5% low gelling temperature agarose. The dorsal half of the cord was removed with a vibratome and immersed in PBS for imaging with 0.3 NA or 0.5 NA objectives (section 2.1). Stacks of 201 slices (1000 ms exposure/slice) were acquired, centered on a superficial layer of neurons in coccygeal segments, at sampling intervals of 649×649×2000 nm/voxel (0.3 NA) and 325×325×1000 nm/voxel (0.5 NA). After removal of dark noise, the stacks were deconvolved with the DeconvolutionLab2 plugin [3] (RL, N = 100) using PSFs computed with the modified G&L model specifically for each imaging configuration. A region of interest around the same group of neurons was extracted from each stack, several consecutive slices averaged and the final image upsampled with bicubic interpolation.

## 3 RESULTS

### 3.1 Adaptation of the Gibson & Lanni model

We begin by using the optical configuration and original notation of Gibson and Lanni [4] (G&L). For simplicity we assume that a coverslip of zero thickness is present (*t*_*g\**_ = *t*_*g*_ = 0). Such virtual coverslip has no impact on optical paths, but maintains a formal separation between the objective and tissue sample compartments (Figure 1). In general, fluorescent sources will be located deep in the specimen compartment and away from the ideal position at the coverslip/specimen interface. In real use the objective front lens, which is immersed in its design medium (*n*_*oil*_ = *n*_*oil*\*_), is moved toward the virtual coverslip to shift the objective’s diffraction focus within the sample. The thickness of the objective compartment is thus reduced from its design value (*t*_*oil*_ ≤ *t*_*oil*\*_) (Figure 1A). If the lens reaches the virtual coverslip (*t*_*oil*_ = 0) we fall in the specific case of an objective directly immersed in the same clearing medium as the specimen (Figure 1B). The optical path difference (OPD) for a point source placed at a distance *t*_*s*_ from the virtual coverslip is then:

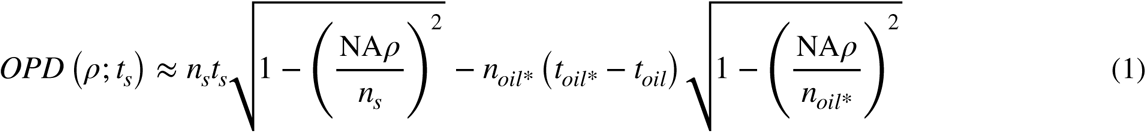

**Figure 1.**
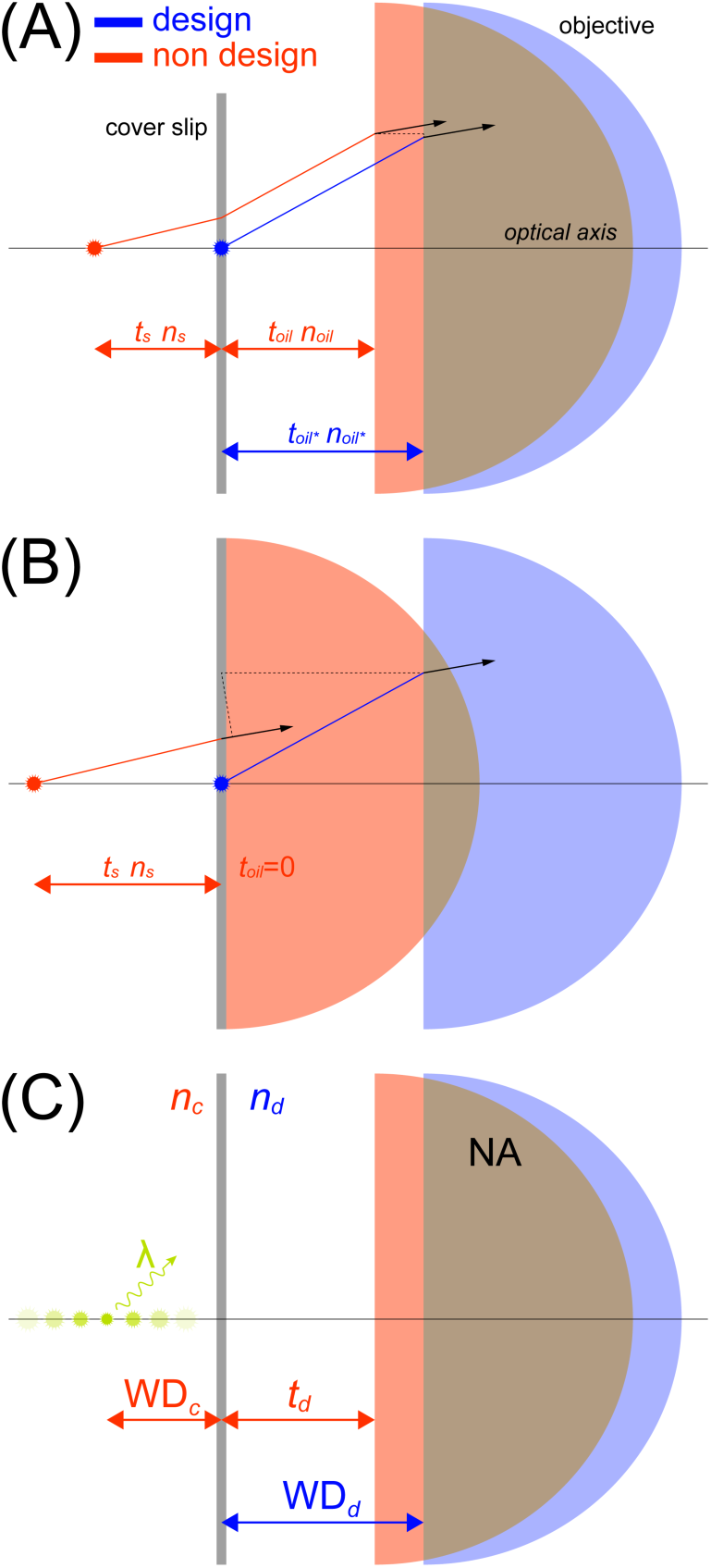
The modelled optical configuration when imaging in mismatched solutions. **(A)** Diagram of the optical configuration modelled in this study, with the notation originally used by G&L [4] and simplified to assume a coverslip of zero thickness. **(B)** This shows the particular case when the coverslip-objective distance is zero, which corresponds to the case of an objective being directly immersed in the tissue clearing medium. **(C)** The same model shown in panel A with our redefined notation and relevant parameters required for evaluating imaging quality using our closed form approximations (see Criteria 1 and 2). NA: numerical aperture of the objective; *n*_*d*_: objective design immersion medium RI; *n*_*c*_: tissue sample clearing medium RI; WD_*d*_: objective working distance in design medium; WD_*c*_: objective working distance in clearing medium; *t*_*d*_: coverslip-objective distance when viewing at the desired depth in the sample (if the objective is directly immersed in the clearing medium this is zero); *λ*: fluorescence emission wavelength.

We now define a new set of variables that are more relevant for our optical configuration:

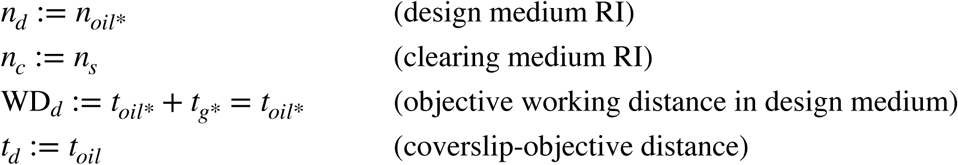

Furthermore, we note that under a paraxial approximation the objective working distance in the clearing medium (WD_*c*_; defined as the distance of the diffraction focus in the specimen compartment from the virtual coverslip; figure 1C) is given by [5]:

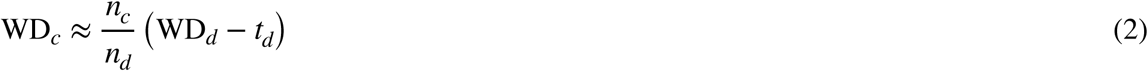

We now define the sum of the right hand side of eq. 2 and an axial displacement variable *z*,as the distance of the point source from the virtual coverslip:

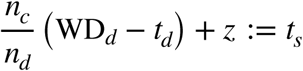

Substituting the newly defined variables in eq. 1 we obtain:

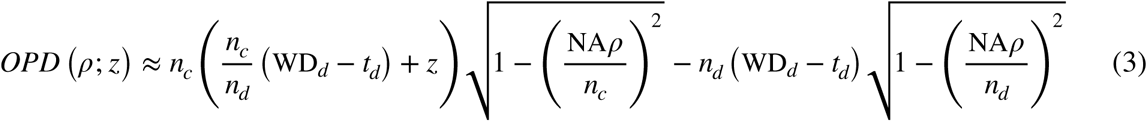

### 3.2 Derivation of approximate expressions for image quality degradation

We estimated the degree of aberration introduced by a mismatch between the clearing and design media, at the approximate diffraction focus (*z* = 0), via Maclaurin expansion in *ρ* of eq. 3:

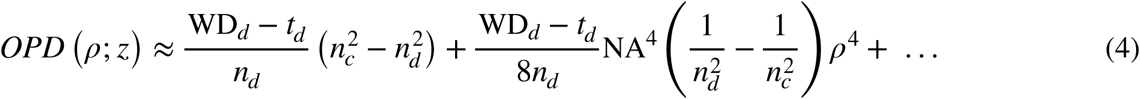

The quartic term in eq. 4 represents primary spherical aberration [18]. Neglecting higher order terms, its coefficient *A*_*s*_ provides a closed-form expression of the amount of aberration affecting the imaging quality of our objective:

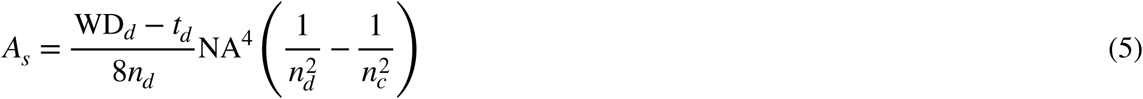

For small differences between the clearing and design medium RIs we have:

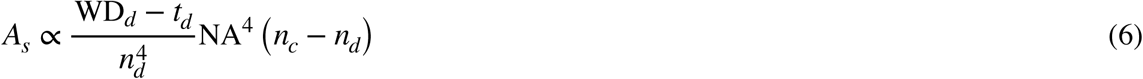

Thus, the magnitude of spherical aberration |*A*_*s*_|:

a. increases as NA^4^, ruling out high aperture objectives unless optimized for the specific clearing medium; even then, small RI mismatches may severely degrade imaging quality;
b. increases as |WD_*d*_ − *t*_*d*_|, suggesting to place the coverslip as close as possible to the tissue to be imaged (*t*_*d*_ ≈ WD_*d*_) or, if directly immersing in the clearing medium (*t*_*d*_ = 0), using objectives with the shortest working distance compatible with the required imaging depth;
c. decreases as 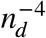, ruling out air objectives except with very low apertures;
d. increases linearly as the RI mismatch |*n*_*c*_ − *n*_*d*_|.

The effect of introducing spherical aberration in a well corrected optical system is to flatten and expand the 3D PSF, mainly along the optical axis (Figure 4 B and C), and redistribute energy to the outer rings of the Airy pattern [7, 8]. This leads to a lower peak brightness of imaged point sources (Figure 2A), a phenomenon quantified by the *Strehl ratio* (the peak brightness of the aberrated PSF divided by that of the unaberrated PSF). In a small aberrations regime this ratio is well approximated by:

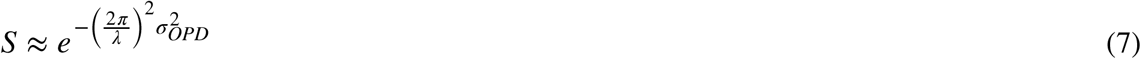

where 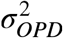 is the variance of the OPD over a circular uniform pupil [19] and *λ* is the fluorescence emission wavelength of the point source. A primary spherical aberration *A*_*s*_ *ρ*^4^ has a standard deviation 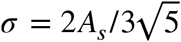. However, in a free focusing system as the fluorescence microscope, the aberration variance can be minimized by introducing a small amount of defocus in a process called *aberration balancing* [20]. Balanced aberration is given by *A*_*s*_ (*ρ*^4^ − *ρ*^2^) and has a smaller standard deviation 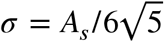, which together with eq. 5 and 7 leads to:

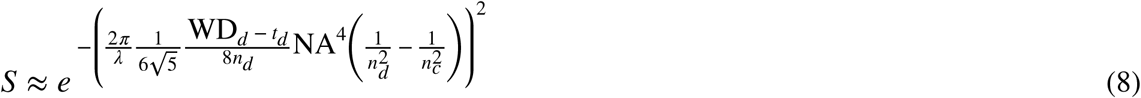

**Figure 2.**
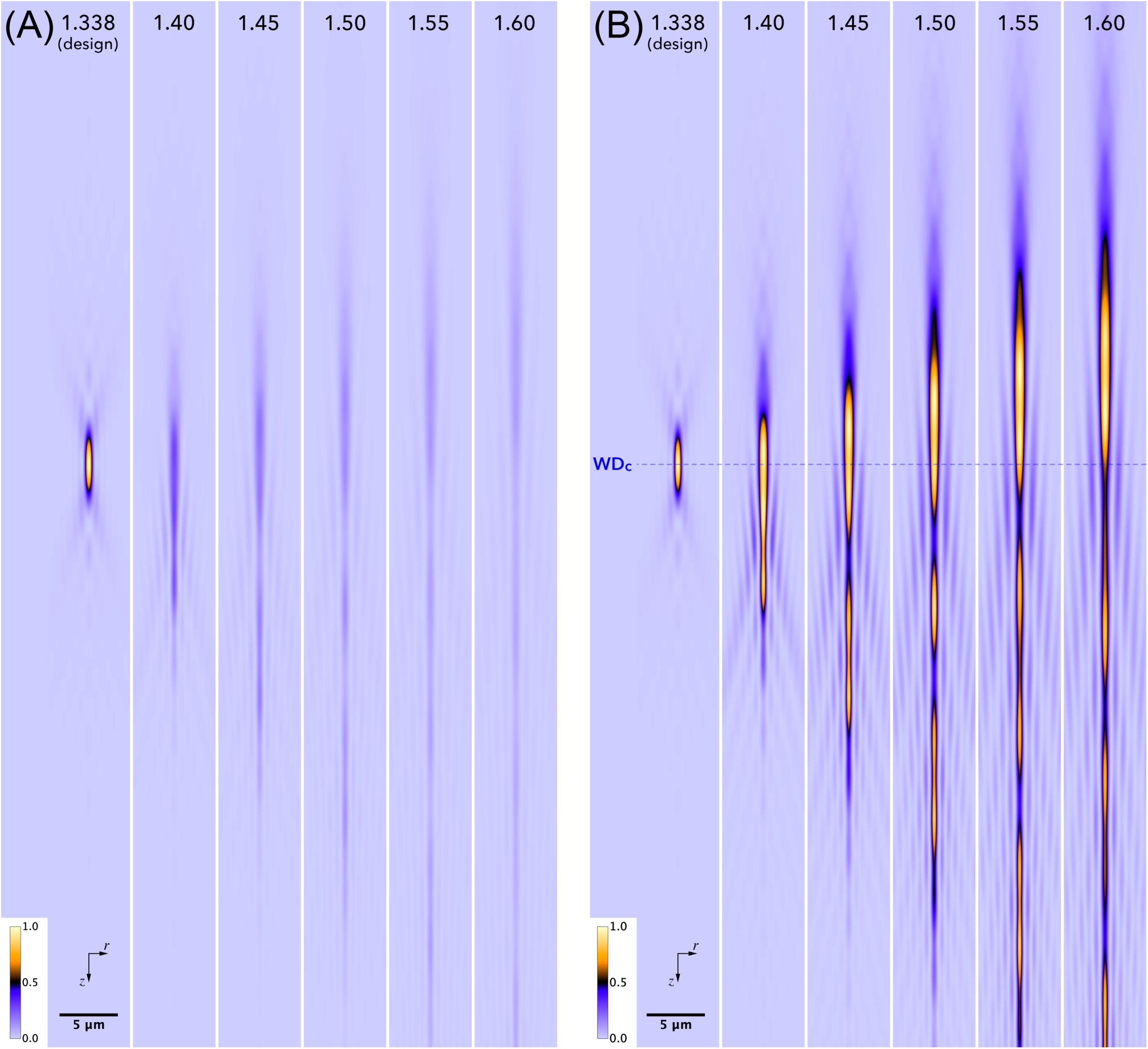
Axial sections of the system’s 3D PSF (from now on simply ‘PSFs’) obtained by integration of the adapted G&L model (eq. 10 and 11) for the extreme case of the objective being directly immersed in the clearing medium (*t*_*d*_ = 0mm); a range of clearing medium refractive indices are explored using a 0.50 NA water immersion objective (20x, WD_*d*_ = 3.50mm). The point source lies on the optical axis at a distance WD_*c*_ + *z* from the objective. **(A)** Aberrated PSFs are normalized to the maximum intensity of the PSF in design conditions, to show their decrease in peak brightness (or Strehl ratio). **(B)** Aberrated PSFs are normalized to their respective maxima to highlight their marked elongation and widening, which leads to a loss of spatial resolution during imaging. The distance of the main peak of the aberrated PSFs from the objective lens (located upwards) is well predicted by eq. 9 (dashed line: WD_*c*_). *λ* = 510nm.

Consistently, when placing the specimen in the objective design medium (*n*_*c*_ = *n*_*d*_) the Strehl ratio is unity, while any deviation (*n*_*c*_ ≠ *n*_*d*_) will decrease its value.

To obtain an improved expression for the objective working distance in the clearing medium, we added the shift caused by the defocus term *A*_*s*_ *ρ*^2^ (see eq. 18 in [21]) to the paraxial approximation of eq. 2:

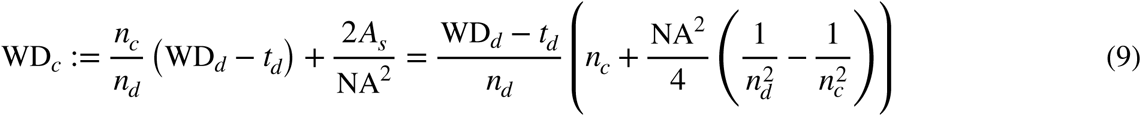

Equation 8 and 9 can be readily applied to any experimental configuration, since their parameters are widely available.

### 3.3 Final PSF model

The final form of our adapted G&L model sees a revised eq. 3 that accounts for the working distance estimated by eq. 9, such that:

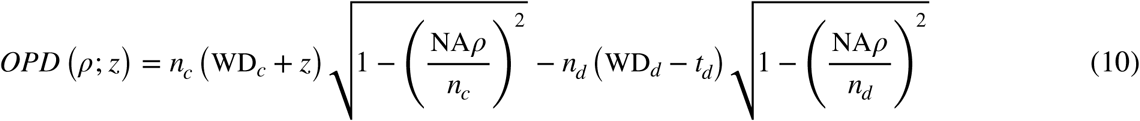

The 2D PSF acquired by the camera sensor when a point source lies on the optical axis at position *z* can be obtained from eq. 5 in [4] by considering that for all practical cases M^2^ ≫ NA^2^ (M is the lateral magnification of the system):

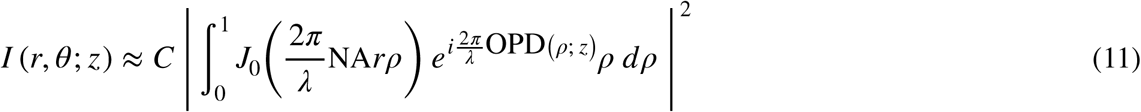

*r* and *θ* are the polar coordinates of a camera pixel back-projected in object space, *C* is a constant, *J*_0_ is the Bessel function of the first kind of order zero. *I* (*r*, *θ*; *z*) defines an infinite set of 2D PSFs as varies, which form the so-called ‘3D PSF’ of the system.

In the following we assessed the predictive power of the the approximate formulae given by eq. 8 and 9, for the particular case of an objective directly immersed in the clearing medium (*t*_*d*_ = 0). This avoided experimental errors associated with measuring the distance between the objective lens and a coverslip while exploring the strongest aberrations attainable with a given objective/clearing medium combination.

### 3.4 The approximate formulae compare favorably with values from computed PSFs

The PSF model described by eq. 10 and 11 was numerically evaluated for *t*_*d*_ = 0 by adapting the open source code of *PSF Generator* [22] for Fiji/ImageJ [13, 14]. We first examined how the Strehl ratio decays when the clearing medium RI departs from the design medium. Figure 2A shows several computed axial sections of the system’s 3D PSF (sections which will be denoted in shorthand as ‘PSFs’), for the moderately severe case of a 0.5 NA long working distance water immersion objective. Intensities were normalized to the maximum value of the unaberrated PSF in water (1.338, design). Figure 3A compares the Strehl ratio approximation given by eq. 8 with values obtained from computed PSFs, for this specific objective and another two from the same product family (0.3 NA and 0.8 NA). As expected, eq. 8 was found to provide a good approximation of Strehl for smaller aberrations (i.e. for smaller NAs and RI mismatches). The dramatic impact of the quartic dependence on NA of the aberrations, indicated by eq. 6, is clear.

**Figure 3.**
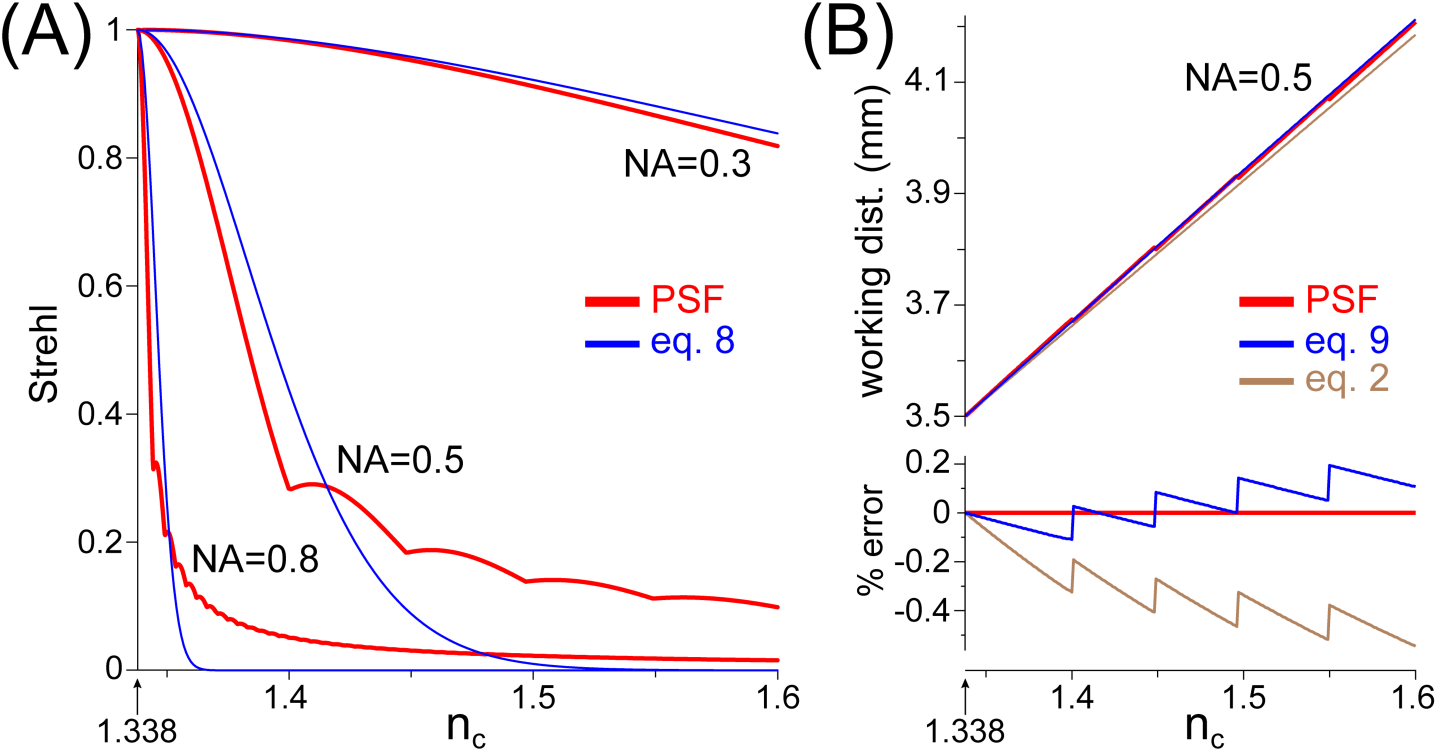
Strehl ratios and working distances predicted by the approximate formulae (eq. 8 and 9), compared to values taken from computed PSFs (adapted G&L model). Here we consider the extreme case of the objective being directly immersed in the clearing medium (*t*_*d*_ = 0mm). **(A)** Strehl ratios as a function of clearing medium RI for three water immersion objectives: 0.30 NA (10x, WD_*d*_ = 3.60mm), 0.50 NA (20x, WD_*d*_ = 3.50mm), 0.80 NA (40x, WD_*d*_ = 3.30mm). Plots show the values from computed PSFs (red) and those predicted by eq. 8 (blue). High apertures objectives are strongly affected by RI mismatches. **(B)** The panel above plots three different working distance estimates for the 0.50 NA objective, as a function of clearing medium RI: distance of the computed PSF principal maximum from the objective front lens (red), the better approximation given by eq. 9 (blue) and that of eq. 2 (brown). The panel below shows the same data as a % error relative to the PSF value. *λ* = 510nm.

We also assessed the approximation of working distance of eq. 9 by using it as the origin of the axial displacement variable *z* in the final PSF model (eq. 10). One would expect the peak of the computed 3D PSF to lie near WD_*c*_ (*z* = 0) irrespective of the degree of aberration. Figure 2B shows the same PSFs of panel 2A but normalized to their respective maxima. Even in the clearing medium with the highest RI (1.60) the PSF maximum is indeed within a few microns of WD_*c*_. Figure 3B compares the predictions made by eq. 9 and 2 with the distance of the computed PSF maximum from the objective lens. The improvement in predictive power offered by eq. 9 over eq. 2 is highlighted in the error graph, which plots their % differences relative to PSF reference value.

### 3.5 Computed PSFs sampled by a synthetic CCD closely match experimental PSFs

To assess whether eq. 10 and 11 generate realistic PSFs, we measured the diffraction pattern of sub-resolution fluorescent sources (section 2.2). An optimal configuration was chosen consisting of the same 0.5 NA water immersion objective and a clearing medium RI of 1.436 (20% FRUIT; [15]). This was predicted to give a PSF with a prominent secondary maximum (Figure 4). Furthermore, the distance between the two peaks was expected to be weakly sensitive to errors in RI (not shown). The model and measured PSFs were similar (Figure 4, main panels), particularly with regards to the distance between the primary and secondary maxima (Figure 4, plot) and the structure of the Airy pattern (Figure 4, below). However, we noted a lower height of the secondary maximum in the measured PSF, as well as a greater lateral elongation near the principal maximum. We hypothesized that these differences could be due to the limited spatial bandwidth of our image acquisition system, particularly in the x-y plane: sampling by the CCD was somewhat below the Nyquist interval. To test this we processed the model PSF by simulating its sampling by the CCD (section 2.4), as proposed by [9]. The resulting PSF was a surprisingly good match to the measured one (Figure 4). Therefore, experimental data provide clear support for the validity of the model under aberrating conditions.

**Figure 4.**
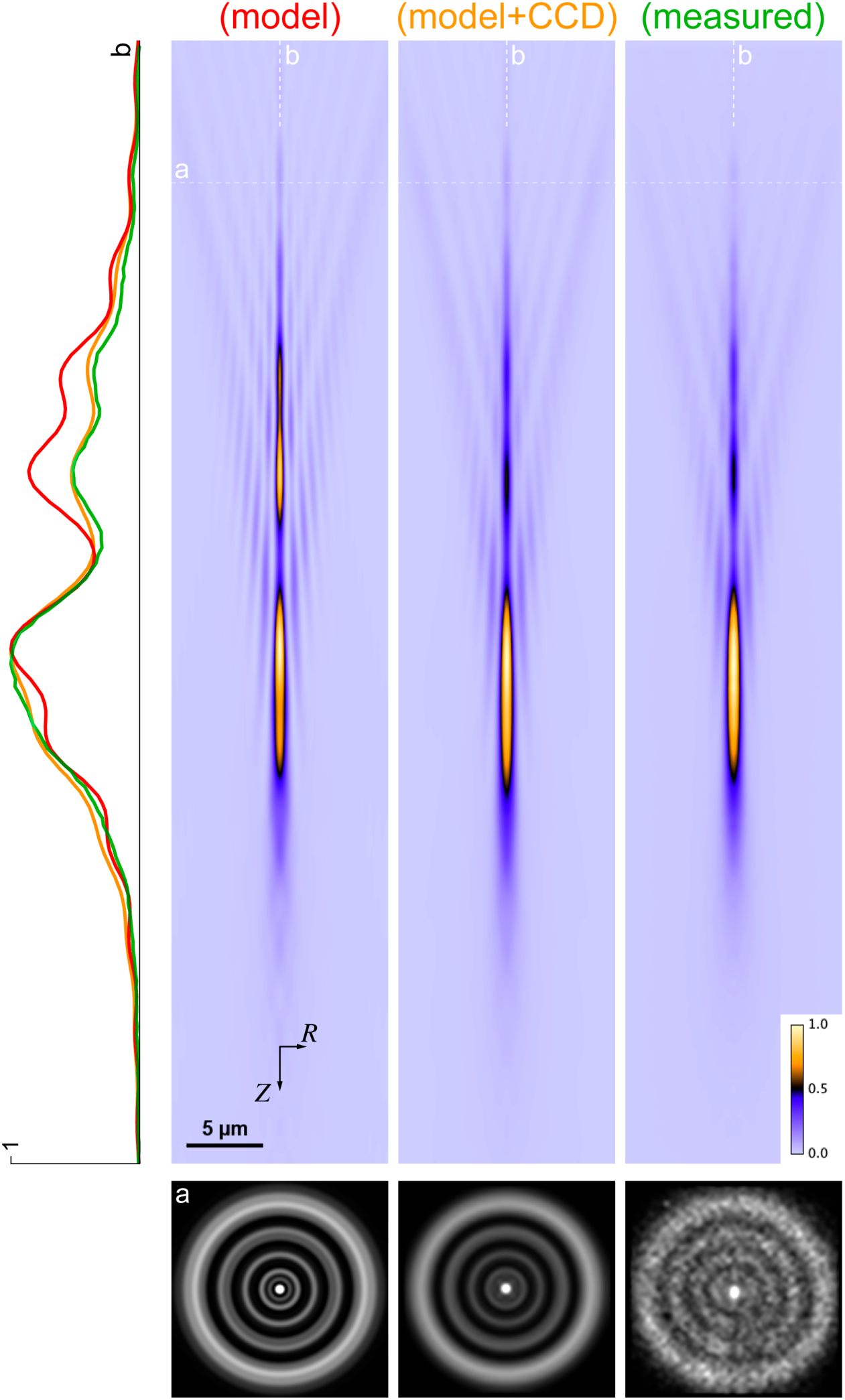
Comparison of the aberrated PSFs predicted by the adapted G&L model, with that measured with sub-resolution fluorescent particles. Here we consider the extreme case of the objective being directly immersed in the clearing medium (*t*_*d*_ = 0nm). **(model)** PSF predicted by eq. 10 and 11 for the same 0.5 NA water immersion objective used in figure 3A assuming a clearing medium RI of 1.436. Here the PSF is displayed in new coordinates (*Z* = − *z*; *R* = *r*) to mimic the common experimental convention where a positive shift of the objective in *Z* brings its diffraction focus deeper in the sample. **(model + CCD)** the model PSF was further processed to simulate the degradation expected to be introduced by the finite size of the pixels in our microscope CCD (sampling errors). **(measured)** experimental PSF obtained with the 0.5 NA objective by averaging stacks from many sub-resolution green fluorescent particles embedded in 1% low gelling temperature agarose and equilibrated with 20% FRUIT clearing solution (measured RI = 1.436). The plot on the left shows the intensity profile of the three PSFs (red: model, orange: model + CCD, green: measured) taken along a central axis (white dashed lines *b*). The Airy patterns below show cross sections of the PSFs along a transverse plane located 30 *µ*m above the point of maximum intensity (white dashed line *a*). In the model *λ* = nm, while for measurements the emission filter pass band was 500 − 520nm.

### 3.6 The Strehl ratio is a good predictor of the decay in spatial resolution due to RI mismatch

Mismatches in RI not only reduce Strehl but also increase the axial and lateral elongation of the 3D PSF (Figure 2). This has a direct negative impact on the spatial resolution of the optical system since it degrades the ability to distinguish two nearby point sources (e.g. Rayleigh criterion). We sought to determine a simple empirical relationship between resolution degradation and RI mismatch. Since the Strehl ratio is well approximated by eq. 8 and it encompasses the aberrations due to RI mismatches, we explored the relationship between axial/radial elongation of the PSF and Strehl.

Due to the complex structure of spherically aberrated PSFs, elongation cannot be reasonably measured in terms of a full-width at half-maximum (FWHM) of the Airy disc. Instead, we adopted a slightly simplified version of the resolution parameter used in [7], with elongation defined by:

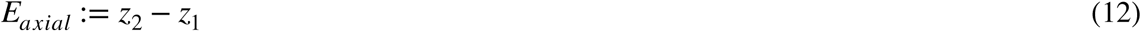

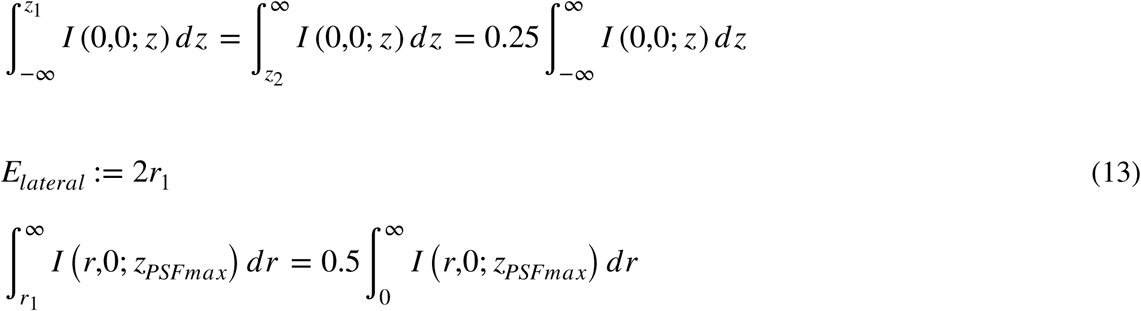

where *z*_*PSFmax*_ is the axial coordinate of the diffraction focus. An example of this approach is shown in graphical form in figure 5A.

We then determined elongation and Strehl on computed PSFs for the three water immersion objectives used in the previous sections, in a range of clearing medium RIs. Figure 5B and C show these data in a compact form as plots of the inverse of elongation normalized to its value in design conditions (i.e. for *n*_*c*_ = *n*_*d*_) versus Strehl. The thin dashed lines represent fits to the data (restricted to Strehl > 0.5) of the function:

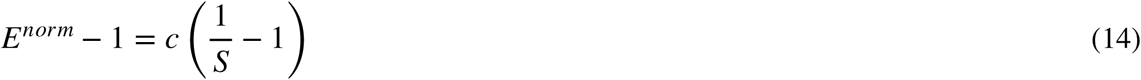

**Figure 5.**
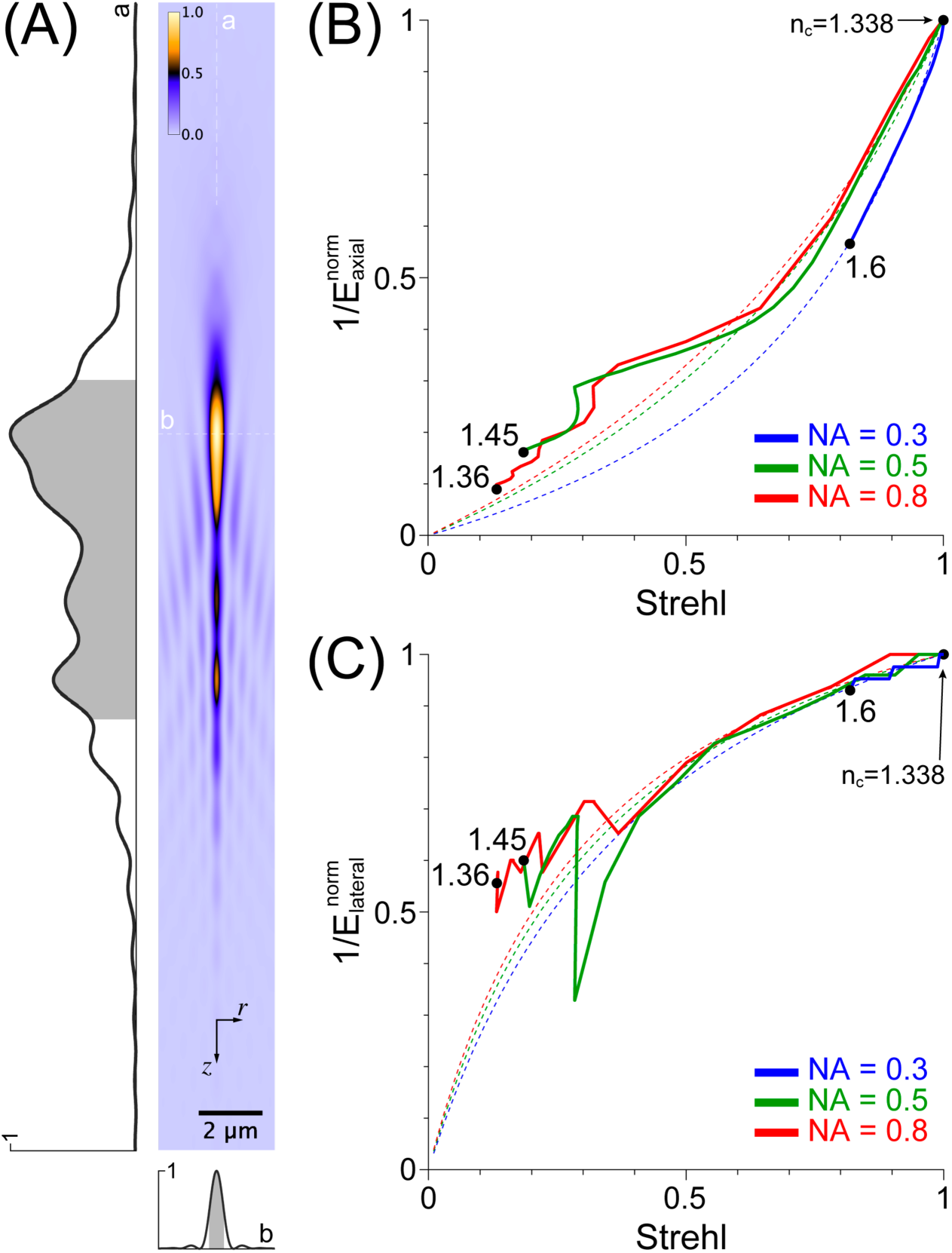
Imaging resolution degrades proportionally to the inverse of the Strehl ratio. Here we consider the extreme case of the objective being directly immersed in the clearing medium (*t*_*d*_ = 0mm). **(A)** The axial and lateral elongation of a system’s 3D PSF determine its two-point discrimination ability (i.e. its spatial resolution). For our aberrated PSFs we determined these two parameters on plots of the intensity along axial or radial lines passing through the diffraction focus (white dashed lines *a*, *b*), as the interval (gray areas) on either side of which (white areas) the integrated intensity was 25% of the total. The PSF shown refers to the same 0.80 NA objective used in figure 3A with *n*_*c*_ = 1.35. **(B)** Plots of the inverse of the axial elongation (normalized to its value in design conditions) versus Strehl ratio for the same three objectives of figure 3A. The clearing medium RIs are shown for the plot endpoints. Dashed lines represent best fits to the data (restricted to a low aberration range of Strehl > 0.5) of a relation of direct proportionality between *normalized axial elongation increase* and *inverse of Strehl increase* (eq. 14) (0.30 NA: c = 3.40, R = 0.997; 0.50 NA: c = 2.29, R = 0.996; 0.80 NA: c = 2.03, R = 0.994). **(C)** Analogous of the plot in B for lateral elongation (0.30 NA: c = 0.32, R = 0.850; 0.50 NA: c = 0.28, R = 0.992; 0.80 NA: c = 0.25, R = 0.987). *λ* = 510nm.

Equation 14, which assumes a simple relationship of direct proportionality between the incremental values beyond unity of normalized elongation and inverse Strehl, gives good results (Figure 5 legend). Two important observations can be made:

a. Axial elongation degrades faster than Strehl (*c* > 1) while the opposite is true for lateral elongation (*c* < 1). This implies that RI mismatches lead to severely reduced optical sectioning capacity before they significantly affect lateral resolution.
b. All three objectives degrade their performance with a similar progression relative to Strehl. Therefore, this parameter can be used as a single synthetic predictor of imaging quality degradation.

Based on these considerations and figure 3A and 5B,C we propose an empirical scale to evaluate whether an objective/clearing medium combination is suitable in terms of brightness and resolution relative to diffraction limited imaging.

#### CRITERION 1

*S* > 0.9 (*excellent*); 0.7 < *S* < 0.9 (*good*); 0.4 < *S* < 0.7 (*mediocre*); *S* < 0.4 (*poor*)

where the Strehl ratio *S* is given by eq. 8 with all parameters (summarized in figure 1C) available to the microscopist. Note that this criterion naturally extends to the general case of the objective and specimen compartments being separated by a coverslip (i.e. *t*_*d*_ > 0).

### 3.7 Better absolute spatial resolution may sometimes be achieved with a lower NA objective

In the previous section we examined the image quality degradation of a given objective when viewing a target located in a non design clearing solution. However, in practice one may be faced with the choice of using one among several available objectives. Given point (ii) in the previous paragraph and the fact that the Strehl ratio decays faster for higher NAs (Figure 3), one can expect that in mismatched media lower NA objectives may achieve smaller axial and lateral elongations than higher NA ones. We verified this seemingly paradoxical behavior both in modeling (Figure 6) and in experiment by imaging the same fluorescent neurons with different objectives (Figure 7).

**Figure 6.**
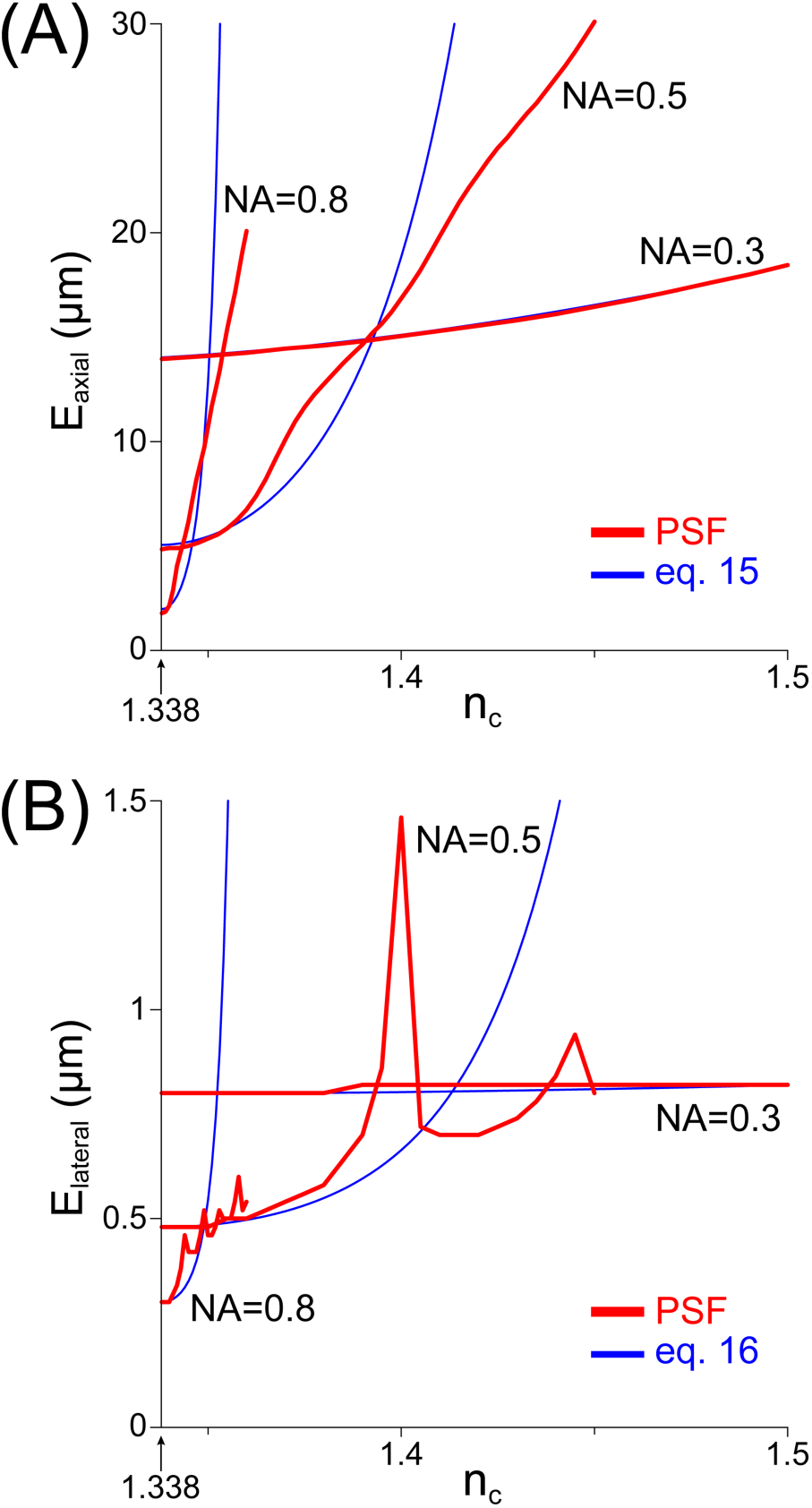
Lower NA objectives may achieve have better resolution than their higher NA counterparts in mismatched media. Here we consider the extreme case of the objective being directly immersed in the clearing medium (*t*_*d*_ = 0mm). Axial (**A**) and lateral elongation (**B**) were determined from computed PSFs for the three water immersion objectives used as test cases in this study. As the RI of the clearing medium departs from the design one, the resolution of higher NA objectives degrades faster until it becomes worse than that of the lower NA ones. Also shown are the approximate elongations predicted by eq. 15 and 16. *λ* = 510nm.

**Figure 7.**
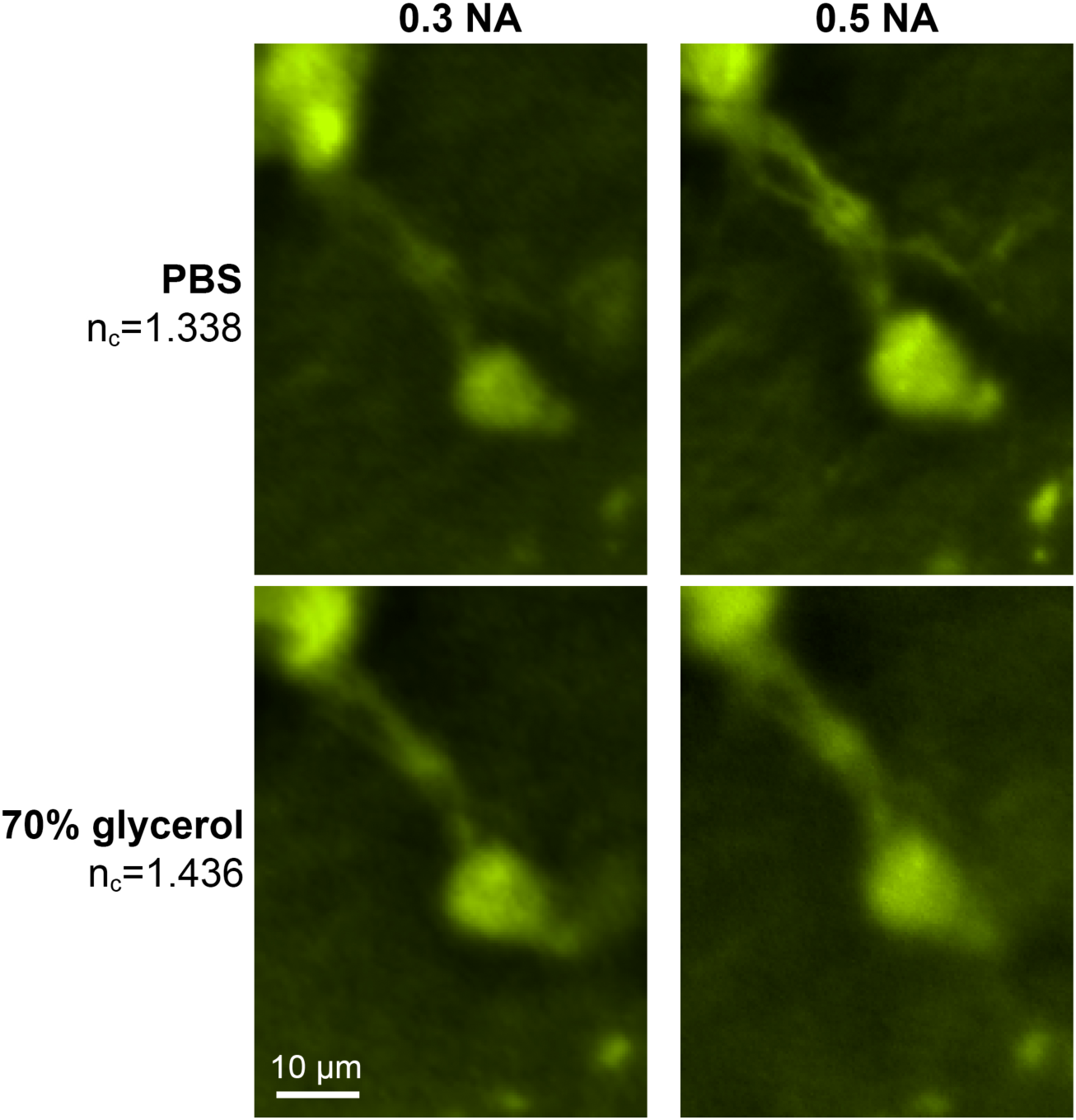
Images of fluorescent neurons, acquired by immersing the objective in a clearing solution of mismatched RI, degrade according to modeling predictions. Fluorescent neurons from the spinal cord of an early postnatal Galanin-eGFP^+/+^ mouse [16] were imaged with 0.3 NA and 0.5 NA water immersion objectives. The cells shown here were located near the cut surface of the horizontally hemisected spinal cord, to allow unobstructed visualization in aqueous (i.e. design) solution when the tissue is opaque. **(Top panels)** tissue was fixed and immersed in design medium (phosphate buffered saline, PBS; *n*_*d*_ = 1.338). As expected from computed PSF elongations (figure 6) and the approximations of eq. 15–16 the higher NA objective was able to resolve finer neuronal processes: 0.3 NA, *E*_*axial*_ = 14*μ* m, *E*_*lateral*_ = 0.8*μ* m; 0.5 NA, *E*_*axial*_ = 5*μ* m, *E*_*lateral*_ = 0.5*μ* m (values from eq. 15–16). **(Bottom panels)** the same tissue and neurons after equilibration with 70% glycerol solution (*n*_*c*_ = 1.436). With the 0.3 NA objective image quality is not significantly degraded, as expected (see figure 3, 5 and 6). In fact, a slight improvement is apparent, which may be attributed to the clearing effect of the high RI solution. With the 0.5 NA objective, however, the switch to glycerol severely affects image quality, bringing it to a lower level than that attained by the 0.3 NA objective under identical conditions, again the expected behavior: 0.3 NA, *E*_*axial*_ = 16*μ* m, *E*_*lateral*_ = 0.8*μ* m; 0.5 NA, *E*_*axial*_ = 68*μ* m, *E*_*lateral*_ = 1.3*μ* m (values from eq. 15–16). All images were obtained by acquiring 3D stacks centered on the neurons and deconvolving them using theoretical PSFs generated with the G&L model.

To obtain closed form approximations of the axial and lateral elongation, we combined known expressions of diffraction limited resolution in fluorescence microscopy [23], with the empirical relationship found in the previous section to obtain:

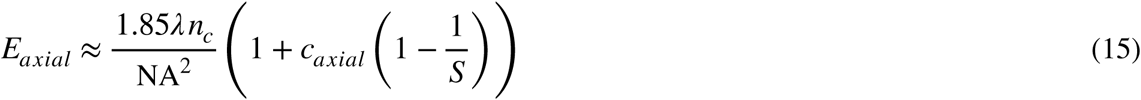

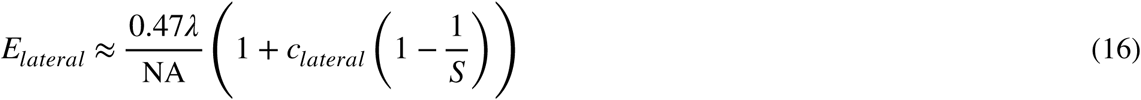

where *S* is given by eq. 8, *c*_*axial*_ = 2 and *c*_*lateral*_ = 0.3 (based on the fits to eq. 14 shown in figure 5B and C). Eq. 15 and 16 were surprisingly effective in predicting the elongations of computed 3D PSFs (Figure 6). Therefore, a gross comparison between several objectives can be made with them.

#### CRITERION 2

*The objective with the lowest axial/lateral elongation as given by eq. 15/16 is best suited for the given clearing medium*.

Ultimately, an accurate prediction of the axial and lateral elongation of an objective’s 3D PSF can be only obtained by numerical integration of the modified G&L model.

## 4 DISCUSSION

An extensive body of work has explored, mostly at the theoretical level, the aberrations introduced by variations in RI along the microscope’s optical path, such as when thin tissue slices are mounted on microscope slides [4–7]. More recently, attention has shifted to methods of correcting such aberrations [8, 24] and to higher-order aberrations due to RI inhomogeneities within biological samples [25]. The current renaissance in the field of tissue clearing, motivated by an interest in viewing deep while preserving fluorescence [2, 26–29], has further aggravated the impact of aberrations due to a combination of high RI clearing solutions and great imaging depth (up to several mm).

While a coverslip is frequently used to image cleared preparations, the direct immersion of the objective in the clearing medium (possibly using an RI-matched coverslip [30]) is an attractive optical configuration: (i) the diffraction focus moves by the same distance as the objective (i.e. axial scaling is unity) and the lateral magnification is the same as in design conditions; (ii) aberrations are depth-of-focus independent and thus a single 3D PSF can be used for deconvolution; (iii) coma-like aberration, which is introduced by even the slightest tilt of a non RI-matched coverslip [31], does not occur; (iv) the entire working distance of the objective (WD_*c*_) can be used; (v) single cell electrophysiology in semi-cleared living tissue may soon become feasible [32]. A disadvantage, however, is that if the objective is not designed for the RI of the clearing medium, the resulting aberrations will be determined by its full working distance (WD_*d*_ in eq. 5) irrespective of the depth of focus in the specimen.

While microscope manufacturers are expanding their catalog of objectives to cover the spectrum of clearing solutions, they are generally very expensive. Even when a nominally optimal objective is at hand, perhaps equipped with a correction collar, one may wish to assess how sensitive imaging quality will be to any residual RI offset. Importantly: (i) the RI of cleared tissue is likely to be somewhat different from that of the clearing solution; (ii) the RI of lab-made clearing solutions is seldom checked with a refractometer. Therefore, it should be of practical interest for the microscopist to rapidly assess *a priori* the imaging performance of a specific experimental configuration. If necessary, one can generally adjust the clearing solution RI without compromising final tissue transparency [15]. The two criteria proposed in this study are simple to calculate: NA, WD_*c*_, *n*_*d*_ are given by the objective manufacturer; *λ* is the centroid of the product of the fluorescent source and emission filter transmittance spectra; *n*_*c*_ is published or can be measured; *t*_*d*_ is the trickiest parameter to determine with precision but is only necessary when using a coverslip.

These criteria were explicitly developed for the widefield fluorescence microscope since the cost of purchasing an optimized objective is more likely to be an issue than for cutting edge microscopes (these are often shared facilities). Furthermore, ongoing advances in deconvolution may greatly improve their computational optical sectioning performance [3, 33]. However, the approximate expressions for Strehl ratio and elongation (eq. 8, 15–16) are relevant also to more advanced microscopes. In the case of the confocal microscope they separately apply to the illumination and detection PSFs [5]. In the light sheet microscope, they apply to the detection PSF [8], while in the two-photon microscope they apply to the ‘single-photon’ illumination PSF [11].

## Acknowledgements

We thank Daniel Sage at EPFL for kindly providing the Java source code of PSF Generator.

